# Nuclear pore density controls heterochromatin reorganization during senescence

**DOI:** 10.1101/374033

**Authors:** Charlene Boumendilrid, Priya Hari, Karl C. F. Olsen, Juan Carlos Acosta, Wendy A. Bickmore

## Abstract

Oncogene induced senescence (OIS) is a cell cycle arrest program triggered by oncogenic signalling. An important characteristic of OIS is activation of the senescence associated secretory phenotype (SASP)^1^ which can reinforce cell cycle arrest, lead to paracrine senescence but also promote tumour progression^2–4^. Concomitant with cell cycle arrest and the SASP activation, OIS cells undergo a striking nuclear chromatin reorganization, with loss of heterochromatin from the nuclear periphery and the appearance of internal senescence-associated heterochromatin foci (SAHF)^5^. The mechanisms by which SAHF are formed, and their role in cell cycle arrest and expression of the SASP, remain poorly understood. Here we show that nuclear pore density increases during OIS and is responsible for SAHF formation. In particular, we show that the nucleoporin TPR is required for both SAHF formation and maintenance. The TPR-induced loss of SAHF does not affect cell cycle arrest but completely abrogates the SASP. Our results uncover a previously unknown role of nuclear pores in heterochromatin reorganization in mammalian nuclei and in senescence, which uncouples the cell cycle arrest from the SASP.

## Main

SAHF formation results from a reorganization of pre-existing heterochromatin – genomic regions decorated with repressive histone marks such as H3K9me3, H3K27me3, MacroH2a and heterochromatin proteins *HP1α,β,γ* - rather than de novo formation of heterochromatin on new genomic regions^5–7^. SAHF appear consecutive to cell cycle arrest and are never observed in replicating cells^5^. Factors implicated in the formation of SAHF include; activation of the pRB pathway,^5^ certain chromatin-associated non-histone proteins^8^, and the histone chaperones HIRA and Asf1a^6,9^. SAHF are proposed to participate in the silencing of pro-mitotic genes, contributing to stable cell cycle arrest in senescence^8^. However, whether SAHF formation is necessary for cell cycle arrest, or for expression of the SASP phenotype, has not been investigated.

In non-senescent cells, a major proportion of heterochromatin is associated with the nuclear lamina (lamina associated domains - LADs), that underlines the nuclear envelope. We hypothesized that a modification of the nuclear envelope could lead to the formation of SAHF, by decreasing the forces localising heterochromatin to the nuclear lamina and/or by increasing factors that repel heterochromatin (Fig. 1a). In agreement with this hypothesis, there is a loss of LADs during OIS^10,11^, with the released heterochromatin then presumably forming SAHF. LaminB1 expression is decreased in OIS and its experimental depletion can facilitate (but is not sufficient for) SAHF formation^10^.

**Figure 1.**
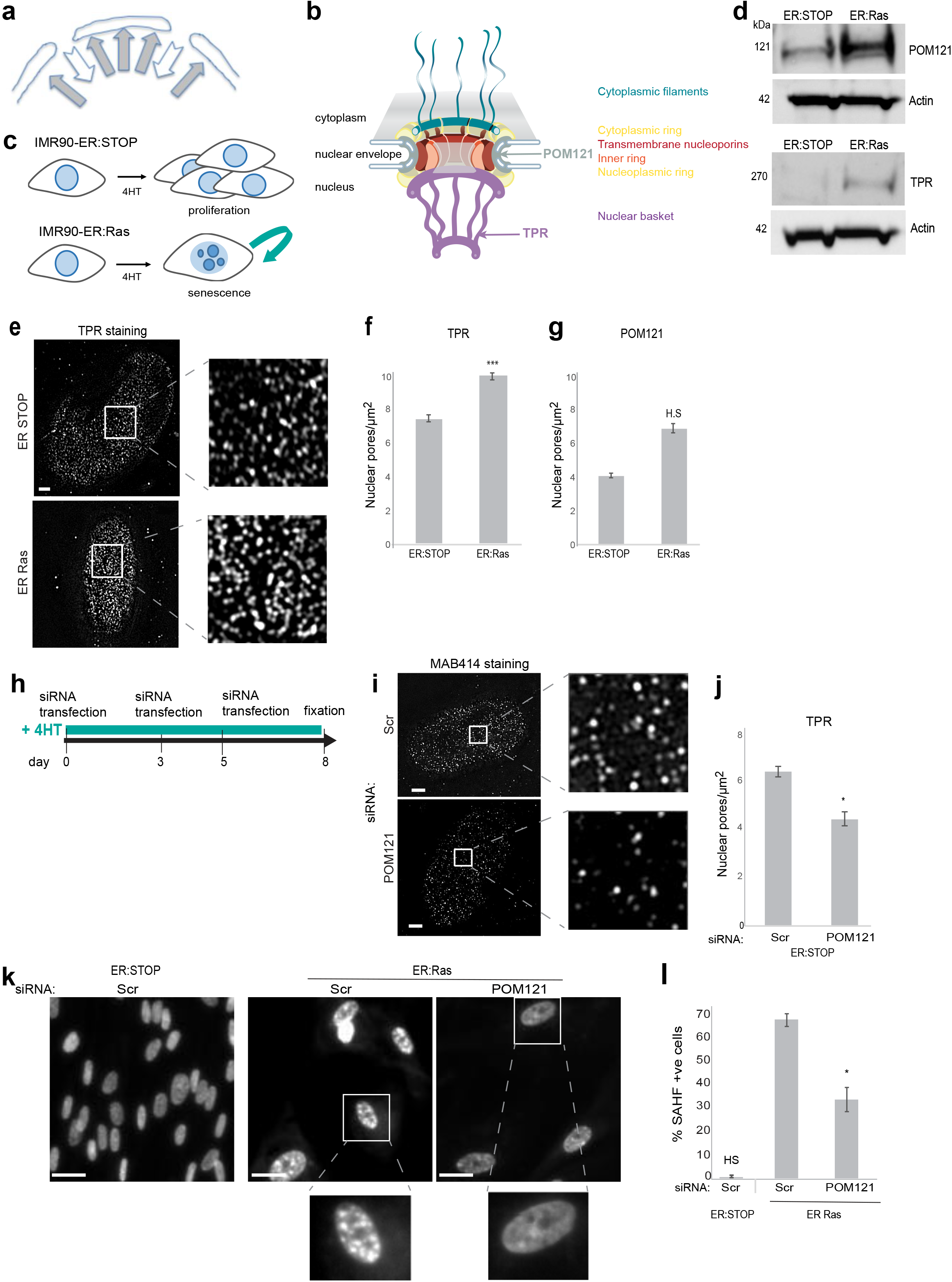
Increase of nuclear pores density in OIS is necessary for SAHF formation. a) Schematic showing the balance of forces attracting heterochromatin to the nuclear lamina and repelling heterochromatin from nuclear pores. b) Model of the nuclear pore complex indicating the position of the nucleoporins TPR and POM121, adapted from ^25^. c) Schematic of OIS induction in IMR90 cells by 4HT and continued proliferation in control ER-Stop cells. d) Western blot showing POM121 (left panel) and TPR (right panel) protein levels in 4HT treated ER-Stop and ER-Ras cells. e) TPR Immunostaining in ER-STOP and ER-Ras cells treated with 4HT. Left: bottom plane of nucleus imaged by SIM. Right: enlargement of the insets. Scale bars 2 μm. f) Mean (+/- SEM) nuclear pore density (pores/μm^2^) in 4HT treated ER-Stop and ER-Ras cells as counted by TPR staining in 3 independent biological replicates, \*\*\**p*=0.0001. g) As for f) but for Pom121 staining. H.S = highly significant *p*=1.3^E-06^. h) Schematic showing depletion experiment as performed for panels J) and K) i) MAB414 (antibody recognizing several nucleoporins) immunostaining in ER STOP cells treated with 4HT after 2 days-knockdown with scramble (Scr) or POM121 siRNAs. Left: bottom plane of nucleus imaged by SIM. Right: enlargement of the insets. Scale bars 2 μm. j) Mean (+/- SEM) nuclear pore density (pores/μm^2^) in 4HT treated ER-Stop cells after knock down with scramble (Scr) or POM121 siRNAs as assayed by TPR staining in 3 independent biological replicates, *= p<0.05. k) DAPI staining of 4HT-treated ER-Stop and ER-Ras cells in controls (Scr) and upon POM121 depletion (siPOM121). Scale bars 10 μm. Bottom: enlargement of the insets. l) Mean (+/- SEM) % of cells containing SAHF in 4HT-treated ER-Stop and ER-Ras cells after knockdown with scramble (Scr) siRNAs and in 4HT-treated ER Ras cells with POM121 SiRNAs. Data from 3 independent experiments. *=p<0.05 HS=highly significant.

Contrasting with studies on the nuclear lamina, the role of nuclear pores in the formation of SAHF has not been studied. Nuclear pores perforate the nuclear membrane, allowing selective import and export of macromolecules into and out of the nucleus. The 120 MDa nuclear pore complex (NPC) is composed of a highly ordered arrangement of nucleoporins (Fig 1b). Whereas most of the nuclear envelope is associated with heterochromatin, the area underneath the nuclear pores is completely devoid of it and NPCs have been proposed to actively exclude heterochromatin^12^. We therefore considered that the relative density of nuclear pores could be important in the balance of forces attracting or repelling heterochromatin at the nuclear periphery and decided to assess the role of NPCs in the formation of SAHF in OIS.

We used a system in which the activity of oncogenic Ras (RAS^G12D^) is induced by addition of 4-hydroxy-tamoxifen (4HT) in human IMR90 cells, leading to OIS - activation of p53 and p16 and expression of SASP proteins3 (Fig. 1c; Extended Data Fig. 1a). Nuclear pores disassemble upon entry into mitosis but are very stable during interphase^13–15^. To stabilise nuclear pore density during differentiation, quiescent cells down-regulate the expression of mRNA encoding nucleoporins^16^. However, RNA expression profiling in OIS cells (ER-Ras) showed that, compared with control ER-STOP (STOP codon) cells, nucleoporin mRNA levels remain unchanged during senescence (Extended Data Fig. 1b). This suggests a potential accumulation of nucleoporins in senescent cells and we confirmed this by immunoblotting for two nucleoporins; POM121 – an integral membrane protein of the NPC central ring^17,18^ and TPR – a large coiled-coil protein of the nuclear basket (Fig. 1b, d). Immunofluorescence and structured illuminated microscopy (SIM)^19^ showed that the increase in nucleoporin levels during OIS results in an increased nuclear pore density (Fig. 1e-g).

As components of the NPC have been shown to interact preferentially with euchromatin^12^, we hypothesized that NPCs may actively exclude heterochromatin and that the density of nuclear pores – i.e. the relative area of the nuclear periphery that repels heterochromatin, may be a critical factor in determining whether heterochromatin is retained at the nuclear periphery or released from it to self-associate in SAHF in the nucleoplasm (Fig. 1a).

To assess whether the increased nuclear pore density we observe in OIS cells is responsible for heterochromatin reorganization into SAHF, we used siRNAs to deplete POM121 (Extended Data Fig. 2a) during the entire course of OIS induction (Fig. 1h). As expected, since POM121 is required for NPC assembly during interphase^13,18^, this led to a decrease in nuclear pore density (Fig. 1i, j and Extended Data Fig. 2b). Consistent with our hypothesis, POM121 depletion resulted in a dramatic reduction of OIS cells containing SAHF (Fig. 1k, l).

**Figure 2.**
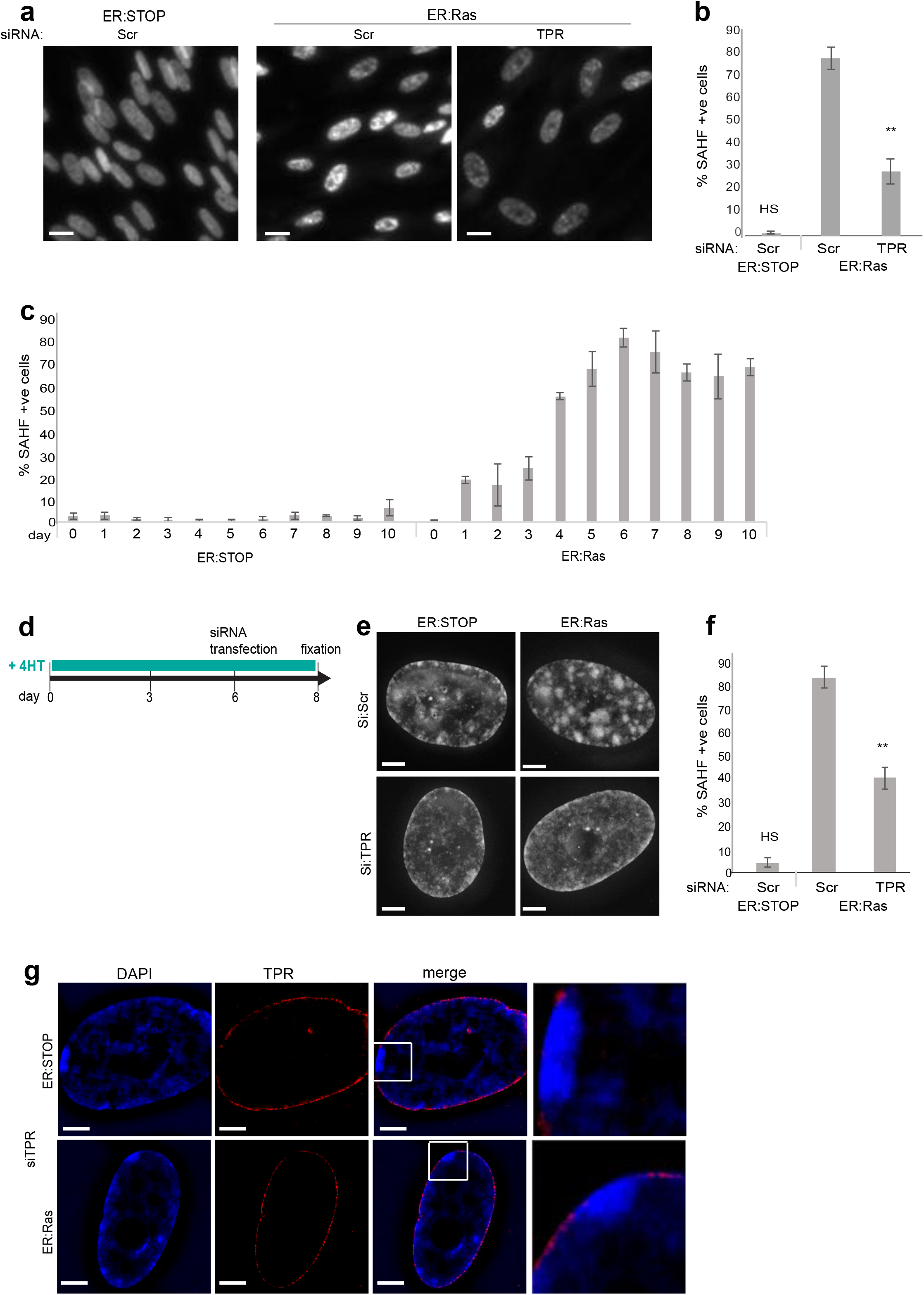
TPR is necessary for SAHF formation and maintenance. a) DAPI staining of non-senescent 4HT treated ER-Stop and OIS (ER-Ras) cells after control scrambled (Scr) SiRNA and upon TPR depletion (siTPR). Scale bars 10 μm. b) Mean (+/- SEM) % of cells containing SAHF in 4HT-treated ER-Stop and ER-Ras cells after knockdown with scramble (Scr) siRNAs and in 4HT-treated ER Ras cells with TPR SiRNAs. Data from 3 independent experiments. **=p<0.01 HS=highly significant. c) Time course of mean (+/- SEM) % cells with SAHF after 4HT-treatment of control (ER-Stop) and OIS cells (ER-Ras). Data from 3 independent experiments. d) Time course for TPR depletion by siRNA late in the OIS programme as performed for panels e-g. e) DAPI staining of 4HT-treated ER STOP and OIS cells ER:Ras in controls (Scr) and upon TPR depletion (siTPR). Scale bars 2 μm. f) Mean (+/- SEM) % of cells containing SAHF in 4HT-treated ER-Stop and ER-Ras cells after knockdown with scramble (Scr) siRNAs and in ER-Ras cells with TPR SiRNAs. Data from 3 independent experiments. **=p<0.01 HS=highly significant. g) DAPI (blue) and TPR (red) staining of 4HT-treated ER-STOP and ER-Ras upon TPR depletion (siTPR) imaged by SIM. Right: enlargement of the insets. Scale bars 2 μm.

TPR has been shown to establish heterochromatin exclusion zones at nuclear pores^20^ and so the increased abundance of TPR at the nuclear periphery of OIS cells, as a result of the elevated nuclear pore density, might be responsible for SAHF formation. We therefore depleted TPR during OIS induction (Extended Data Fig. 3a, b). This did not affect nuclear pore density (Extended Data Fig. 3c). However similarly to POM121 depletion, TPR depletion led to the loss of SAHF (Fig. 2a, b). To rule out off target effects, we confirmed these results with four independent siRNAs targeting TPR (Extended Data Fig. 3d-f). We conclude that TPR is necessary for the formation of SAHF.

**Figure 3.**
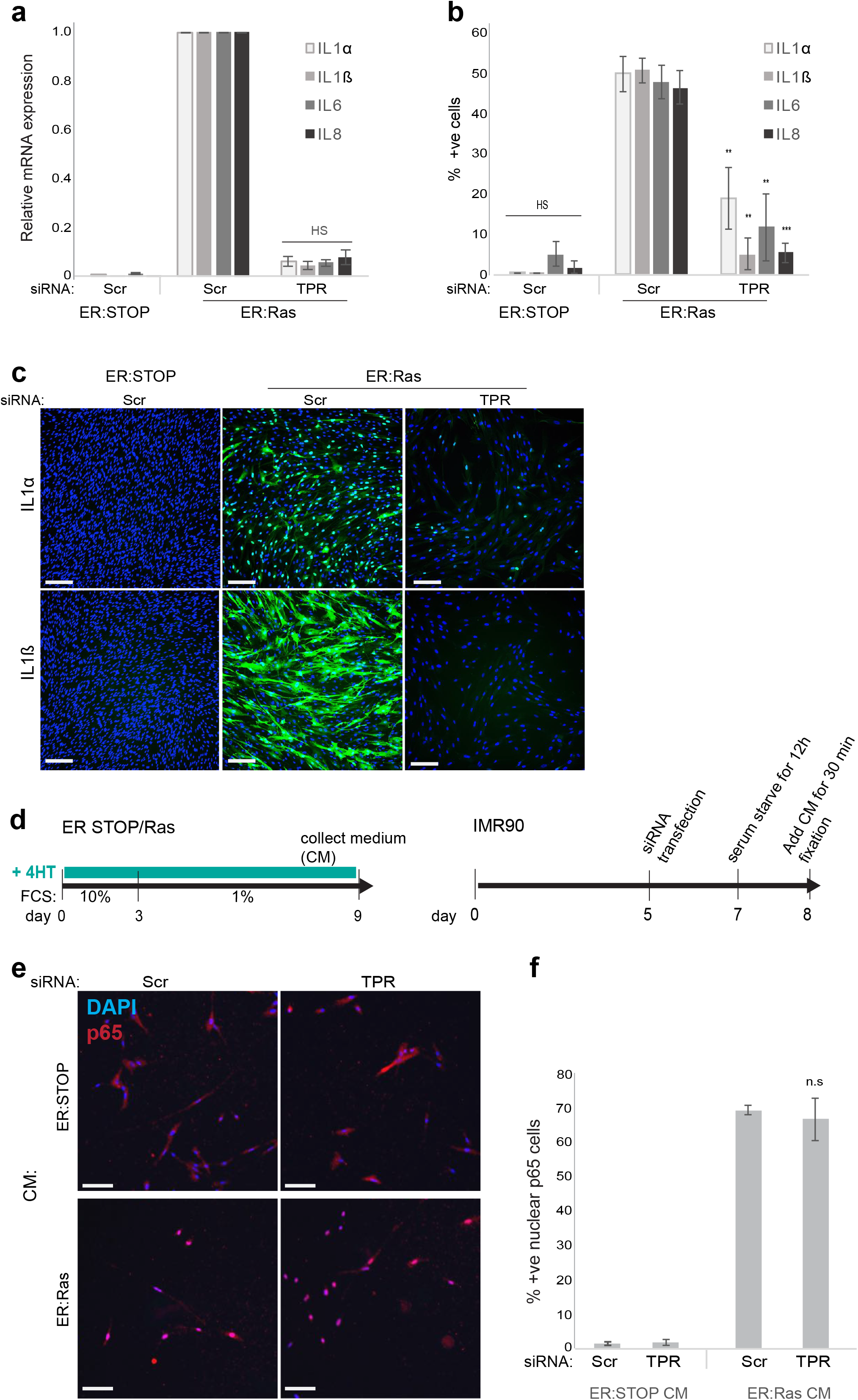
TPR is necessary for the SASP. a) Mean (± SEM) mRNA level, measured by qRT-PCR for SASP genes (IL1α, IL1β, IL6,IL8), in 4HT-treated ER-Stop and ER-Ras cells after knockdown with scramble (Scr) siRNAs and in 4HT-treated ER-Ras cells with TPR SiRNAs. Expression is shown relative to ER-Ras cells transfected with Scr siRNAs. Data from 3 independent experiments. b) Mean (± SEM) % of cells positive for SASP cytokines (IL1α, IL1β, IL6,IL8) in 4HT-treated ER-Stop and ER-Ras cells after knockdown with scramble (Scr) siRNAs and in 4HT-treated ER-Ras cells with TPR SiRNAs, measured by immunostaining. Data from 3 independent experiments. **=p<0.01 ***= p<0.001, HS=highly significant. c) Immunostaining (green) for SASP cytokines IL1α and IL1β in DAPI (blue) stained nuclei of 4HT-treated ER-Stop and ER-Ras cells subjected to RNAi with scrambled (Scr) siRNAs or siRNAs targeting TPR. Scale bars 100 μm. d) Schematic of NFκb nuclear import assessment as performed for panels e and f. Medium (CM) of ER STOP and Ras cells was collected after 9 days of culture in presence of 4HT. CM was then added to IMR90 cells upon scramble (Scr) or TPR depletion by SiRNA. NFκb-p65 import in the nucleus of IMR90 cells was assessed 30 min after addition of CM. e) Immunostaining (red) for p65 in DAPI (blue) stained cells for scrambled (Scr) or TPR (siTPR) depleted IMR90 cells upon addition of collected medium (CM) from 4HT-treated ER-Stop and ER-Ras cells. Scale bars 100μm. f) Mean (+/- SEM) % of cells containing nuclear p65 staining in scrambled (Scr) or TPR (siTPR) depleted IMR90 cells upon addition of collected medium (CM) from 4HT-treated ER-Stop and ER-Ras cells. Data from 3 independent experiments. n.s=non significant.

To assess whether TPR was necessary for the maintenance as well as the formation of SAHF, we performed a time course experiment to determine when SAHF are formed after 4HT addition. The percentage of cells containing SAHF increased from 1 day after 4HT treatment of ER-Ras cells, reaching a maximum at 6 days (Fig. 2c). We therefore depleted TPR with siRNAs 6 days upon 4HT addition, when SAHF have already formed (Fig. 2d). We observed a dramatic reduction of cells containing SAHF two days later (day 8) (Fig. 2e, f). TPR staining revealed that our siRNA depletion under these conditions was only partial and we could observe loss of SAHF in cells specifically depleted for TPR, whereas SAHF were maintained in cells where knockdown was incomplete (Extended Data Fig. 3g). Closer observation revealed that in cells with partial depletion of TPR there was a relocalization of heterochromatin towards the nuclear periphery in patches that correspond to sites of TPR-depletion (Fig. 2g). We conclude that exclusion of heterochromatin from the nuclear periphery by TPR is necessary for both the formation and maintenance of SAHF during OIS.

We next wanted to investigate the phenotypic consequences of SAHF loss in OIS ER-Ras cells depleted for TPR. OIS cells depleted for TPR did not show any defect in cell-cycle arrest as assayed by BrdU incorporation (Extended Data Fig. 4a, b) and there was proper activation of p16, p21 and p53 cell cycle regulators (Extended Data Fig. 4c). This suggests that SAHF are dispensable for cell-cycle arrest. However, in the absence of SAHF after TPR depletion, we observed a complete loss of SASP as exemplified by lack of IL1α, IL1β, IL6 and IL8 mRNA (Fig. 3a) and protein induction (Fig 3b, c, Extended Data Fig. 4d). This result was confirmed with four individual TPR siRNAs (Extended Data Fig. 4e). To rule out that SAHF and SASP loss upon TPR depletion was due to a general defect in nuclear transport, we assessed NFκB import into the nucleus upon induction of paracrine senescence^2,3,21,22^ (Fig. 3d). We observed no difference in NFκB import in cells depleted with TPR or scrambled siRNAs (Fig. 3e, f), suggesting that TPR depletion does not lead to nuclear protein transport defects that could explain SAHF and SASP loss.

**Figure 4.**
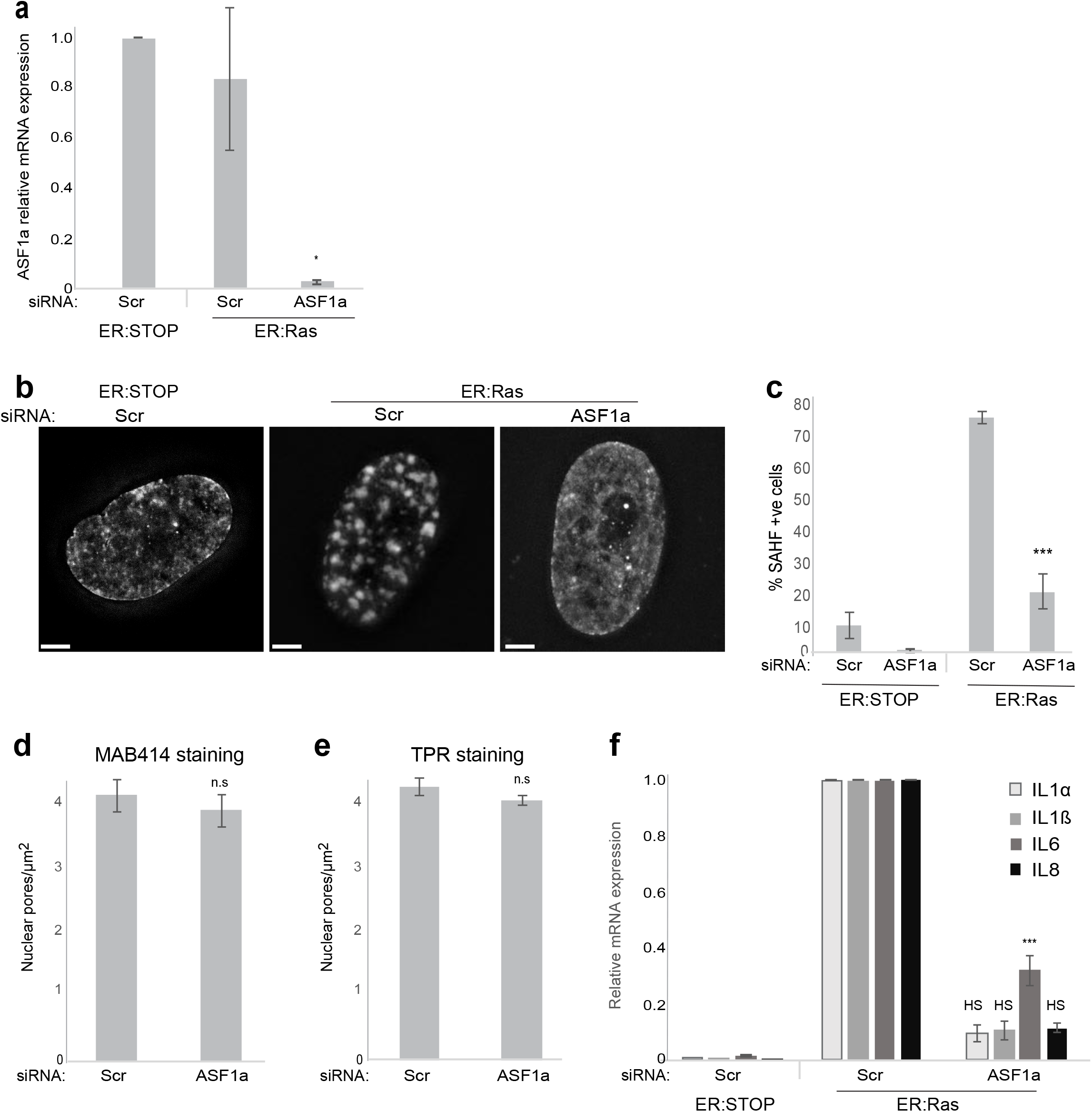
SAHF formation is necessary for the SASP. a) Mean (± SEM) ASF1a mRNA level, established by qRT-PCR, in 4HT-treated ER-STOP and ER-Ras cells after knockdown with scramble (Scr) or ASF1a siRNAs. Expression is shown relative to ER-STOP cells transfected with Scr siRNAs. Data from 3 independent experiments. *= p<0.05. b) DAPI staining of 4HT-treated ER-STOP and ER-Ras cells in controls (Scr) and upon ASF1a depletion (siASF1a). Scale bars 2 μm c) Mean (+/- SEM) % of cells containing SAHF in 4HT-treated ER-Stop and ER-Ras cells after knockdown with scramble (Scr) siRNAs and in ER-Ras cells with ASF1a SiRNAs. Data from 3 independent experiments. ***=p<0.001. d) Mean (+/- SEM) nuclear pore density (pores/μm^2^) in 4HT treated ER-Stop cells after knock down with scramble (Scr) or ASF1a (siASF1a) siRNAs as counted by MAB414 staining in 3 independent biological replicates, n.s= non significant. e) As in d) but for TPR staining. n.s= non significant. f) Mean (± SEM) mRNA levels, measured by qRT-PCR, for SASP genes (IL1α, IL1β, IL6, IL8), in 4HT-treated ER-Stop and ER-Ras cells after knockdown with scramble (Scr) siRNAs and in 4HT-treated ER-Ras cells with ASF1a SiRNAs. Expression is shown relative to ER-Ras cells transfected with Scr siRNAs. Data from 3 independent experiments. ***=p<0.001, HS=highly significant.

Similarly to some other nucleoporins, TPR has been shown to be present in the nucleoplasm as well as at nuclear pores^23^. To assess whether it is the increase in nuclear pore density – and consequent increased TPR abundance at the nuclear periphery - in OIS that is necessary for SASP or whether TPR has an independent role in the SASP, we assessed SASP upon POM121 depletion. POM121 is only present within the NPC. We confirmed that decreased nuclear pore density upon POM121 depletion did not affect cell-cycle arrest (Extended Data Fig. 5a), but the SASP was impaired (Extended DataFig. 5b-d).

Our results suggest that heterochromatin reorganization into SAHF is necessary for SASP during OIS. To further support this hypothesis, and to rule out the possibility that nuclear pores regulate SASP through another independent mechanism, we used a different mean to deplete SAHF in ER Ras cells. The histone chaperone ASF1a has been shown to be required for SAHF formation^6,9^ and indeed its depletion (Fig. 4a) led to a loss of SAHF in ER-Ras cells, (Fig. 4b, c), similarly to what we had observed in TPR or POM121 depleted cells^6,9^. ASF1a depletion does not affect nuclear pore density (Fig. 4d, e), but as for TPR and POM121 depletion, there is a dramatic loss of SASP upon ASF1a depletion in ER Ras cells (Fig. 4f), further confirming that SAHF are necessary for SASP.

Our data demonstrate that an increase in nuclear pore density is responsible for the eviction of heterochromatin from the nuclear periphery by TPR and the consequent formation of SAHF in OIS. Similar mechanisms could be conserved in other type of senescence as nuclear pore density is also increased in replicative senescence^24^. The organisation of chromatin relative to the nuclear periphery has generally been considered from the point of view of interactions between (hetero)chromatin and components of the nuclear lamina. Here we demonstrate that the repulsion of heterochromatin by nuclear pores is another important principle of nuclear organisation and it will be interesting to establish whether the modulation of nuclear pore density also influences the 3D organisation of the genome during development.

## Methods

### Cell culture

Cells culture were cultured in standard conditions, 37°C, 5% CO2, in Dulbecco’s Modified Eagle Medium (DMEM) supplemented with 10% fetal bovine serum (FBS). IMR90 cells were obtained from ATCC. IMR90 ER: RAS and ER:Stop cells were produced by retroviral infection of IMR90 human diploid fibroblast cells with pLNC-ER:RAS and pLXS-ER:Stop retroviral vectors respectively as described^3^. To induce Ras translocation in the nucleus 4-hydroxy-tamoxifen (Sigma) diluted in DMSO was added to cells to a concentration of 100 nM. During senescence induction 4HT containing-medium was changed every 3 days.

### SiRNA transfection

2×10^5^ IMR90, ER-STOP and ER-Ras cells were transfected using Dharmafect transfection reagent (Dharmacon) with a 30 nM final concentration of siRNAs. Predesigned siRNAs (Dharmacon) were used to knock down gene expression as follows:

Scr: on target plus non targeting control pool D-001810-10-59 TPR: on target plus smart pool L-010548-00

TPR-6: on target plus J-010548-06

TPR-7: on target plus J-010548-07

TPR-8: on target plus J-010548-08

TPR-9: on target plus J-010548-09

POM121: on target plus smart pool L-017575-04

ASF1a: on target plus smart pool L-020222-02-0020

### mRNA expression analysis

mRNA expression was determined by IonTorrent mRNA sequencing using the Ion AmpliSeq™ Transcriptome Human Gene Expression Kit. 6 independent biological replicates were analysed and adjusted p-value were calculated by Benjamini and Hochberg (BH) and FDR multiple test correction. Data analysis was performed using Babelomics-5(http://babelomics.bioinfo.cipf.es).

### Immunoblotting

1×10^6^ cells were lysed in RIPA buffer and protein concentration was determined using the Pierce BCA protein analysis kit. 15 μg of proteins were loaded and ran into NuPage 3-8% Tris acetate gels (Invitrogen). Transfer of proteins into a nitrocellulose membrane was performed using the iBlot 2 gel transfer device (Thermofisher). Immunoblotting was done using the following antibodies at the indicated dilutions:

TPR: Abcam, ab84516 (1:200)

Pom121: Millipore AB6041 (1:500)

Actin: Santa Cruz Biotechnology, sc-1616 (1:2000)

### Immunofluorescence

2×10^5^ cells were seeded on coverslips and grown during the course of senescence induction. Cells were fixed in 4% paraformaldehyde (pFa) for 10 min at room temperature, permeabilized in 0.1% Triton X100 for 10 min, blocked in 1% BSA for 30 min, incubated with primary antibodies diluted in 1% BSA for 1h and with fluorescently labelled secondary antibodies (Life Technologies) for 45 min. Coverslips were counterstained with DAPI and mounted in Vectashield (Vectorlabs).

### Imaging by Structured Illumination Microscopy (SIM) and measurement of nuclear pores density

The bottom plane of cells was imaged by 3D SIM (Nikon N-SIM) and reconstructed using NIS element software after immunofluorescence with following antibodies at the indicated diutions:

Pom121 (Millipore AB6041) (1:250)

TPR (Abcam, ab84516) (1:500)

MAB414 (Abcam, ab24609) (1:50)

15 nuclei were imaged for each condition and 5 ROI of 100×100 pixels were analyzed/nucleus. Individual nuclear pore complexes in each ROI were counted manually.

### Measurement of SAHF positive cells

Cells were stained with DAPI and observed using epifluorescence microscopy. 100-200 cells were counted per condition and the percentage of SAHF positive cells was calculated.

### BrdU incorporation

Cells in culture were incubated with 10 μM 5-Bromo-2’ – deoxyuridine BrdU (Sigma) for 16 hours prior to fixation. Immunofluorescence staining was conducted as described in the immunofluorescence section and detected using the BrdU Antibody (BD Pharmingene 555627) in the presence of 1 mM MgCl_2_ and 0.5 U/ul DNaseI (Sigma D4527).

### β-Galactosidase staining

SA-β-Gal staining solution was prepared using 20x KC(100 mM K_3_FE (CN)_6_ and 100 mM K_4_Fe (CN) _6_*3H_2_O in PBS), 20x X-Gal solution (ThermoFisher Scientific) diluted to 1x in PBS/1 mM MgCl_2_ at pH 5.5-6. Staining was conducted overnight on cells fixed with glutaraldehyde.

### Detecting SASP, tumour suppressors and BrdU by high content microscopy

Detection of SASP proteins, tumour suppressors and BrdU positive cells by high content microscopy is described at : https://doi-org.ezproxy.is.ed.ac.uk/10.1007/978-1-4939-6670-7_9

### Quantitative reverse transcriptase PCR

Total RNA was extracted using the RNeasy minikit (QIAGEN). Complementary DNAs were generated using Superscript II (Life technologies). PCR reactions were performed on a Lightcycler 480 (Roche) using SYBR Green PCR Master Mix (Roche). Expression was normalized to β-actin. Sequences for the primers used are as follows:

Pom121-Fw TTCAACGTGAGCAGCACAAC

Pom121-Rev CAAAAGTGTTGCCGAAAGGTG

TPR-Fw CTGAAGCAATTCATTCGCCG

TPR-Rev GGCATATCTTCAGGTGGCCC

ASF1a-Fw CAGATGCAGATGCAGTAGGC

ASF1a-Rev CCTGGGATTAGATGCCAAAA

Actin-Fw CATGTACGTTGCTATCCAGGC

Actin-Rev CTCCTTAATGTCACGCACGAT

IL1α-Fw AGTGCTGCTGAAGGAGATGCCTGA

IL1α-Rev CCCCTGCCAAGCACACCCAGTA

IL1α-Fw TGCACGCTCCGGGACTCACA

IL1β-Rev CATGGAGAACACCACTTGTTGCTCC

IL6-Fw CCAGGAGCCCAGCTATGAAC

IL6-Rev CCCAGGGAGAAGGCAACTG

IL8-Fw GAGTGGACCACACTGCGCCA

IL8-Rev TCCACAACCCTCTGCACCCAGT

### Statistics

All experiments were performed in a minimum of 3 biological replicates. Error bars are standard error of the mean. P-values were obtained by two sample equal variance, 2 tails t-test.

## Acknowledgments

C.B. was supported by a H2020 Marie-Curie IF (655350 – NPCChr) and by a prize for young researchers from the Bettencourt-Schueller foundation. JCA is supported by a Career Development Fellowship from Cancer Research UK. W.A.B is supported by a Medical Research Council (MRC) University Unit grant MC_PC_U127527202. We thank Robert Illingworth (MRC HGU, Edinburgh) for the Volcano plot of mRNA expression.

## Authors contribution

CB and WAB conceived the experiments and designed the experiments together with JCA. P.H performed qRT-PCRs and immunostaining for cytokines of the SASP and siRNA transfections for many of the experiments. CB conducted most of the other experiments including; super-resolution microscopy, counting of nuclear pore densities and identification of cells containing SAHFs. KCFO assisted with immunoblotting. CB and WAB wrote the manuscript with input from all authors.

## Competing interests

The authors declare that they have no competing interests.

## Materials and correspondence

Correspondence and material requests should be addressed to WAB

